# Identification and removal of contamination in palaeoproteomic analysis of dental enamel

**DOI:** 10.1101/2025.05.30.657004

**Authors:** Zandra Fagernäs, Sofie Sieling, Gaudry Troché, Jan K. Bakker, Jesper V. Olsen, Frido Welker

## Abstract

Analysis of archaeological and palaeontological dental enamel allows for the study of taxonomy, phylogenetic relationships, and genetic sex of Pleistocene fauna and hominin remains. However, high-quality data is required for the study of degraded, chemically modified ancient proteomes. Protein contamination may hinder acquisition of such data, due to the abundance and better preservation of modern contaminating proteins. Here, we artificially contaminate a Pleistocene woolly rhinoceros tooth in order to investigate decontamination of archaeological dental enamel. We find that although the contaminating proteins are not identified, likely due to a lack of denaturation and digestion steps in archaeological dental enamel protein extraction protocols, contamination leads to significant loss of endogenous proteomic information. We thereafter compare five published decontamination methods, and find that a simple water or bleach wash prior to demineralization is most efficient at removing protein contamination. The water wash does not affect the endogenous proteome, while a bleach wash may lead to a loss of shorter intercrystalline peptides. We therefore conclude that decontamination is necessary in the study of ancient dental enamel, to ensure maximal retrieval of endogenous proteomic information, and can easily be achieved by incorporating a water or bleach wash step into the protein extraction protocol.

## Introduction

Palaeoproteomic analyses of archaeological and palaeontological dental enamel have significantly grown in numbers over the past few years. Given that dental enamel is the hardest tissue in the vertebrate body, consisting primarily of hydroxyapatite, it is favorable in terms of protein preservation^1^. Enamel proteins are therefore among the oldest uncontestedly recovered proteins, with peptides surviving for millions of years^2–6^. The dental enamel proteome is highly conserved^7^, but still contains enough amino acid variation to have been applied to phylogenetic analyses of extinct taxa^2,5,8^. Additionally, genetic sex determination is possible through the enamel protein amelogenin, which is expressed on the sex chromosomes, leading to two forms (amelogenin X and amelogenin Y) being expressed, which have amino acid sequence differences^9^. Through palaeoproteomic analysis of enamel, it is thereby possible to both learn about taxonomic identity and evolutionary relationships, as well as genetic sex and therefore sexual dimorphism, enabling applications to deep-time and more recent (osteo)archaeological analysis.

Analysis of archaeological dental enamel proteomes is, however, not straightforward, as the ancient proteomes are degraded and modified over time. The proteins are fragmented and chemically modified^10^, to a point where they may be completely lost from the archaeological record^11^. Additionally, modern contamination is introduced into the ancient proteome from a wide range of sources, such as the burial environment, handling and storage^12^. For Pleistocene bone, it has been shown that decontamination prior to protein extraction is necessary, as the proteome composition may otherwise be altered, or low-abundance proteins and protein regions may be lost^13^. As many of the dental enamel proteins are not expressed outside the tissue, researchers have thus far often assumed that it is not necessary to include a decontamination step into protein extraction protocols, given that bioinformatic separation of enamel-derived and contaminant proteins is relatively straightforward. Further, the extraction protocols used for archaeological enamel proteins commonly do not include reduction, alkylation, or enzymatic digestion^1^, and it is thereby presumed that modern proteins would not be detected through mass spectrometry. However, the impacts of protein contamination on ancient dental enamel proteomes have not been experimentally studied.

Despite the above arguments, some palaeoproteomic publications do include a claimed decontamination step in their dental enamel proteome extraction protocols. When a decontamination step is included, there is currently no consensus in the field of which method is most efficient, with approaches ranging from mechanical surface removal^14,15^, to treatment with hydrochloric acid^16,17^ or hydrogen peroxide^18,19^, as well as more complex approaches combining several decontamination steps^14,20^. The efficiency of such decontamination approaches, or the effect they have on endogenous enamel proteins, have thus far not been quantified.

Here, we explore the effects of modern protein contamination on an archaeological enamel proteome through artificial contamination of a Pleistocene woolly rhinoceros tooth. Modern dog saliva was chosen for the artificial contamination, given the previous success of this approach^13^, and the approximation of realistic long-term contamination from various sources through this large and complex proteome. We find that although the salivary proteins cannot be detected through typical palaeoproteomic analyses, the effects of this contamination on the endogenous enamel proteome are significant. There is loss of information ranging from fewer identified MS2 spectra, to a decrease in the number of proteins and peptides identified, resulting in the absence of low-abundance protein regions in final protein sequence reconstructions. The recovered proteome is also differing in terms of peptide length, hydrophobicity, and post-translational modifications after contamination. These results clearly show that contamination is a concern for palaeoproteomic dental enamel analysis, and we therefore compare different decontamination approaches. We find that the most efficient approach, returning the proteome nearly to its original state, is a wash with molecular grade water or bleach (sodium hypochlorite) prior to demineralization. A water wash has no impact on the endogenous dental enamel proteome, whereas a bleach wash may decrease recovery of shorter intercrystalline peptides. As this is a simple and straightforward step to incorporate into protein extraction protocols, we recommend that decontamination is conducted prior to analysis of ancient dental enamel, in order to achieve maximal information content from each destructive sampling.

## Results

### Contamination

First, we investigated if the artificial contamination can be identified through commonly employed palaeoproteomic methods for the study of dental enamel. Using a database consisting of enamel, bone, and salivary proteins from modern rhinoceros and dog proteomes, an unspecific search was conducted with MaxQuant^21^, with settings following what is commonly used for studies of ancient dental enamel. However, no proteins were identified as unambiguously stemming from saliva. This is not surprising, since the modern dog proteins introduced during the contamination event are complete, undegraded proteins^13^. Dental enamel protein extraction protocols generally do not include denaturation or digestion steps, as the dental enamel proteins are already fragmented *in vivo*. As a result, dog proteins are likely present as more intact proteins within our extracts, and will generally not be identifiable through the employed extraction and analysis approach.

Next, we compared the identified proteomes between the original (uncontaminated) and contaminated enamel, in order to explore any effects on the endogenous proteome of the thus far undetected contamination. The number of spectra that are acquired during mass spectrometry determine how much data there is to analyse downstream. A higher number of MS spectra was on average acquired from original (6,223.06±17.06; mean ± 1 standard deviation) than contaminated (5,357.57±554.80) samples, although the difference is not statistically significant (ANOVA, F=7.294, p=0.054). The number of acquired MS2 spectra does not significantly differ between the original and contaminated samples either (ANOVA, F=3.749, p=0.125). However, the number of identified MS2 spectra does significantly differ (ANOVA, F=14.78, p=0.0184), as does the percentage of identified MS2 spectra (ANOVA, F=12.57, p=0.0239; Figure 1a), with the contaminated samples having lower values in both cases. This indicates that although the contamination may not alter the number of obtained raw spectra, it does decrease our ability to identify them.

**Figure 1.**
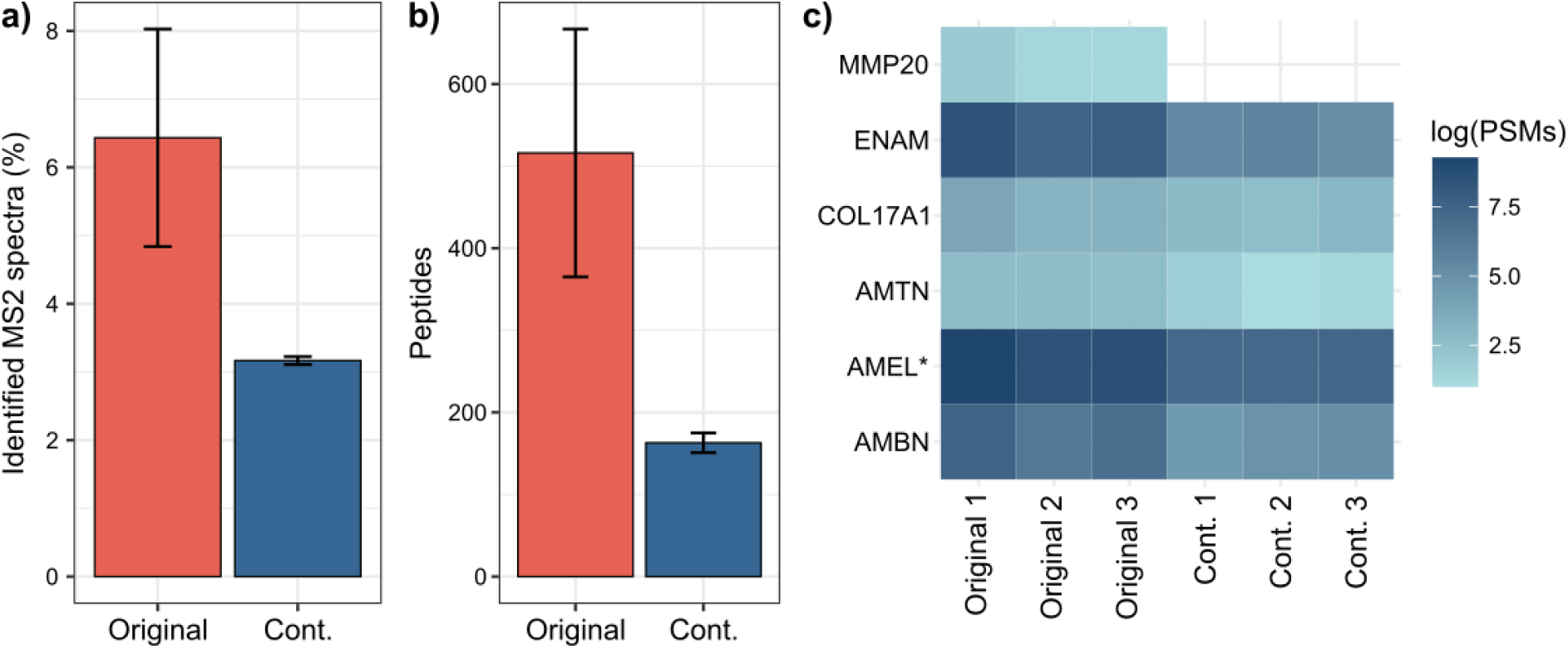
Loss of proteomic information after artificial contamination. a) Percentage of identified MS2 spectra, b) Number of identified dental enamel peptides, c) Total PSM count (log-scale) for enamel-specific proteins. Bars in a-b show the mean of the replicates (n=3) with error bars showing ±1 SD. Cont. = Contaminated.

Looking closer at which proteins were present, for both original and contaminated samples, 20-28 protein groups were identified. A large number of these are, however, different collagen proteins, indicating that traces of dentine were present on the enamel. In all analyses hereafter, the dataset is filtered to only contain dental enamel proteins^1,5^, as they are the focus of this study, but excluding albumin and alpha-2-HS-glycoprotein, as they are ubiquitously expressed across different tissues. This filtered dataset contains 8.61±0.13 dental enamel protein groups for the original samples, and 6.30±0.32 for the contaminated samples, with a significant difference between the sample groups (ANOVA, F=133.7, p=0.0003). The number of identified peptides is also significantly different between the groups (ANOVA, F=16.31, p=0.0156), with 516.04±150.98 peptides for the original samples and 162.95±12.00 peptides after contamination (Figure 1b). Additionally, the number of dental enamel PSMs (peptide spectrum matches) differs significantly between original and contaminated extracts (ANOVA, F=13.56, p=0.0212), with 693.67±225.52 PSMs for the original samples and 211.00±26.15 for the contaminated samples. There is thereby significant loss of proteomic data after a sample has been contaminated, even if the contamination itself cannot be identified.

It is apparent that proteomic information is lost due to contamination, so we investigated whether this loss of information is random or non-random across protein groups. One of the dental enamel proteins, MMP20 (Matrix metalloproteinase-20) could not be identified in any of the contaminated samples, but was present in all original samples (Figure 1c). MMP20 is the least abundant protein in the original samples, indicating that its disappearance may be due to low-abundance peptides being masked by contamination. By inspecting coverage across proteins, it is apparent that there is an overall loss of intensity after contamination. This leads to low-intensity protein regions commonly being completely lost. For example, for amelogenin a region around amino acid position 80-100 is lost in all contaminated samples, and a region around position 140-160 is lost in two out of three contaminated extracts (Figure 2a). A similar pattern can also be seen for ameloblastin (AMBN; Figure 2b) and enamelin (ENAM, Figure 2c).

**Figure 2.**
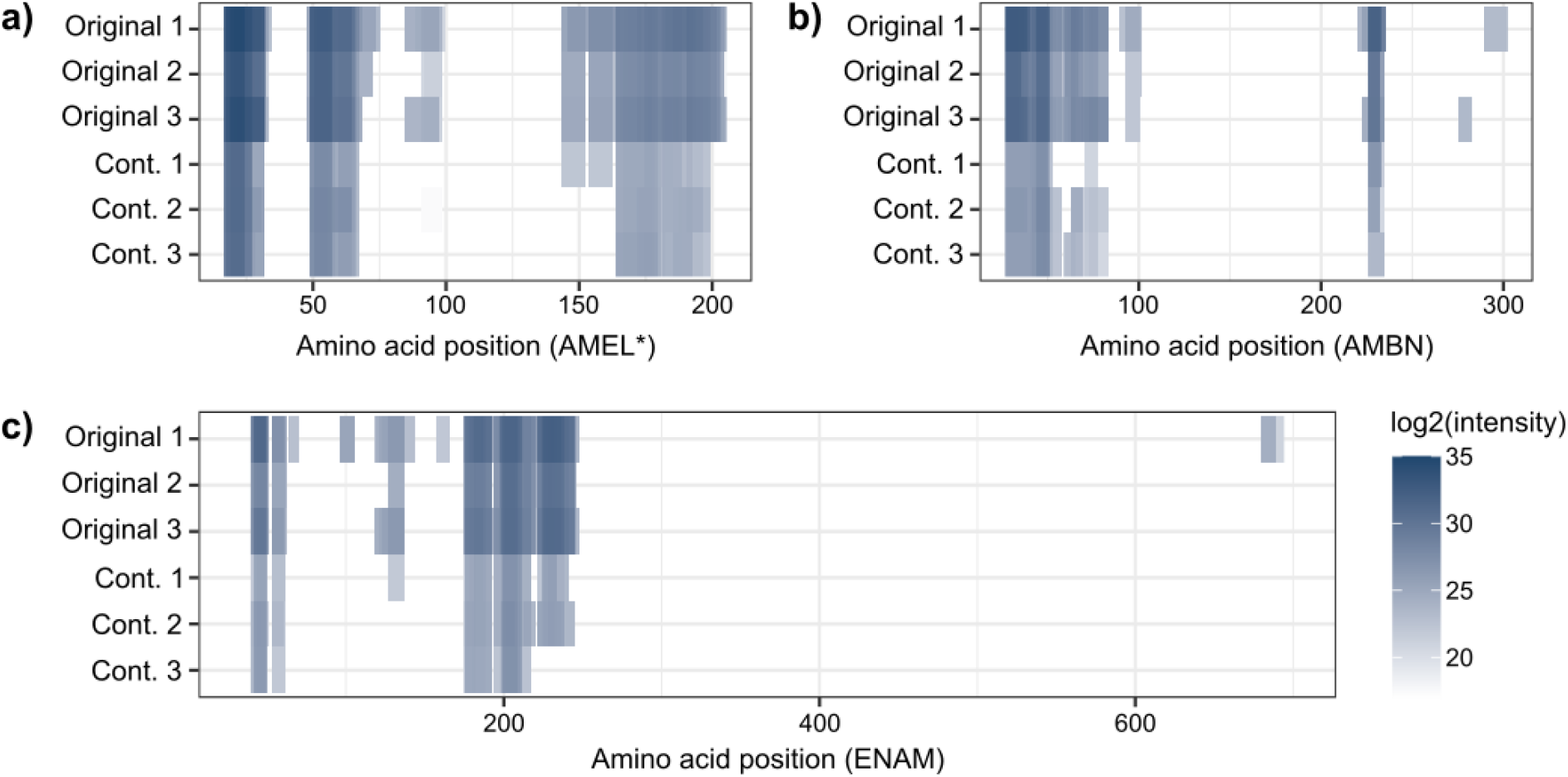
Coverage of enamel proteins. a) Amelogenin, based on amelogenin X (AMEL*), b) Ameloblastin (AMBN), and c) Enamelin (ENAM). Cont. = Contaminated.

Comparing peptide and amino acid properties between the original and contaminated samples may indicate if there is a bias in which type of peptides are lost after contamination. We find that mean enamel peptide length is not different between the contaminated and original samples (ANOVA, F=0.918, p=0.392). Mean hydrophobicity of peptides is, however, different between the original and contaminated samples (ANOVA, F=35.1, p=0.0041), with the peptides in the contaminated samples being on average less hydrophilic (−0.523±0.037) than peptides present in the original samples (−0.698±0.035). The fraction of phosphorylated serines (S; weighted by peptide intensity) differs significantly between original and contaminated samples (ANOVA, F=13.72, p=0.0208), being higher in the contaminated samples (0.491±0.074) than the original samples (0.315±0.036). Deamidation of asparagine and glutamine (N and Q) does not differ between original and contaminated samples (ANOVA, F=0.224, p=0.66 and F=4.212, p=0.109, respectively). Modern protein contamination, while undetected, thereby influences several key parameters of the recovered ancient dental enamel proteome. However, deamidation, which is a commonly used metric for assessing relative antiquity of a proteome, may not be able to detect contamination of the dental enamel proteome.

### Decontamination

As the results above show, contamination can cause significant loss of proteomic information, even in cases when this contamination is not detectable through the methods that are commonly used in palaeoproteomics. It is therefore necessary to perform decontamination prior to protein extraction from dental enamel, to ensure that the maximal amount of endogenous information can be retrieved from each destructive sampling. We compare five methods of decontamination (wash with sodium hypochlorite (bleach), hydrochloric acid (HCl), ethylenediaminetetraacetic acid (EDTA), or molecular grade water, as well as UV irradiation) on subsamples from the same contaminated enamel specimen as analysed above. We evaluate the decontamination methods using metrics that were above found to differentiate the original and contaminated samples, focusing on only enamel-specific proteins (except for analyses of identified MS2 spectra).

The number of identified MS2 spectra varies significantly between the decontamination methods (ANOVA, F=7.017, p=0.0013), with specific differences being identified between original-bleach, original-EDTA, and original-no decontamination (Tukey’s HSD, p<0.05 in each case). Similarly, the percent of identified MS2 spectra also differs significantly between methods (ANOVA, F=4.197, p=0.0127), with significant differences between original-EDTA and original-no decontamination (Tukey’s HSD, p<0.05 in each case). This indicates that most decontamination methods have a variable effect, as they do not significantly improve information recovery compared to the fully contaminated samples. However, EDTA and (partly) bleach washes are significantly different from the original samples. It should, however, be noted that bleach treatment is known to remove intercrystalline enamel proteins ^22^, which may be causing the reduced number of identified MS2 spectra, especially considering a difference cannot be detected between bleach treatment and the original samples when considering the percentage of identified spectra.

The number of recovered enamel proteins significantly depends on the decontamination method (ANOVA, F=6.836, p=0.0019) with specific differences between original-UV, original-EDTA, original-HCl and original-no decontamination, as well as no decontamination-water (Tukey’s HSD, p<0.05 in each case). The number of recovered peptides also depends of the method (ANOVA, F=6.031, p=0.0033), specifically between original-EDTA and original-no decontamination, as well as between water-no decontamination (Tukey’s HSD, p<0.05 in each case). Finally, the number of enamel PSMs depends on decontamination method (ANOVA, F=5.362, p=0.0055; Figure 3a) with significant differences between original-EDTA and original-no decontamination, as well as water-no decontamination (Tukey’s HSD, p<0.05 in each case). For all the above metrics, there are significant differences between a water wash and no decontamination, but not between the original samples and a water wash, indicating that a water wash is efficiently removing contamination.

**Figure 3.**
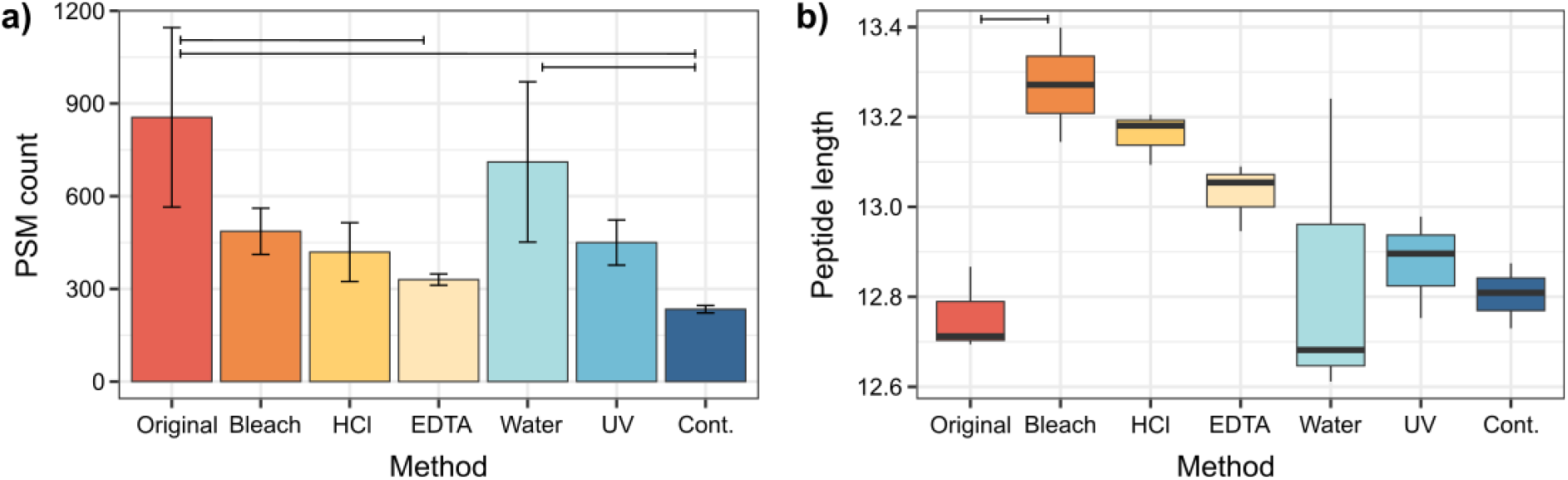
Efficiency of different decontamination methods in recovering the original proteome. a) Number of identified enamel PSMs, and b) Mean peptide length. Horizontal lines indicate statistically significant differences between methods. Cont. = Contaminated.

The overall intensity of peptides of AMEL*, AMBN and ENAM are improved by most decontamination methods, although none of the decontamination methods lead to the same regions being recovered as in the original samples (Figure 4a-c). A water wash will in general lead to most of the same regions being recovered as from the original samples, and may potentially even recover additional regions (e.g., around position 100-110 in ABMN; Figure 4b). A bleach wash also largely recovers the same protein regions as in the original samples, including some regions in AMBN and ENAM not recovered by a water wash; however, we also note that the first region commonly recovered for ENAM by all extraction approaches is not covered after a bleach wash (Figure 4c). For the other decontamination approaches, HCL, EDTA and UV, regions with sequence coverage are largely similar to the non-decontaminated extracts.

**Figure 4.**
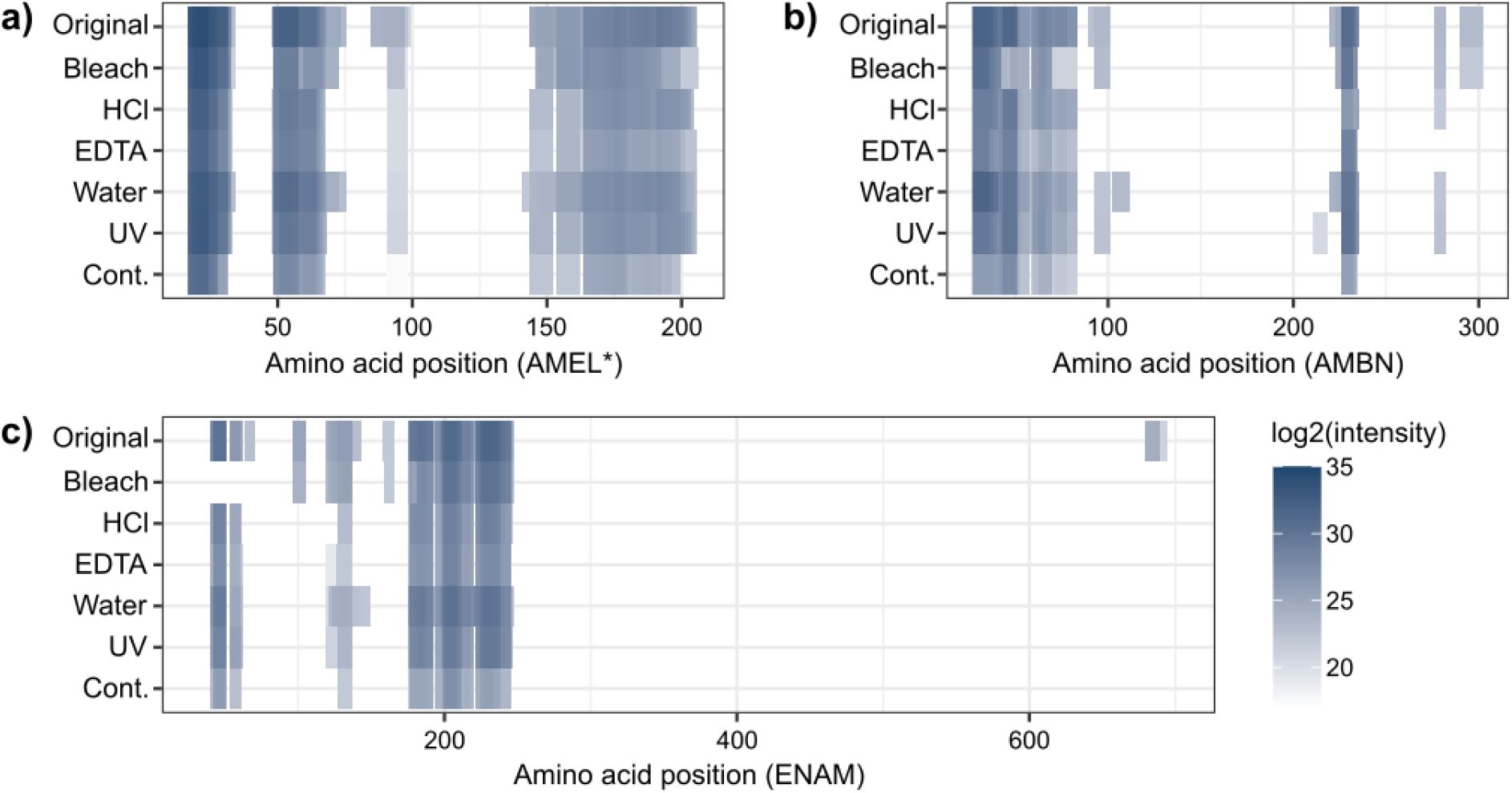
Coverage of enamel proteins after different decontamination methods. a) Amelogenin, based on amelogenin X (AMEL*), b) Ameloblastin (AMBN), and c) Enamelin (ENAM). Intensities are the mean of all replicates for each method. Cont. = Contaminated.

Focusing on the number of covered amino acid positions (Figure 5), a significant difference is present between the decontamination methods for ameloblastin (ANOVA, F=3.582, p=0.0255), with a significant difference only between original-no decontamination (Tukey’s HSD, p=0.0313). For amelogenin, the methods also differ significantly (ANOVA, F=5.174, p=0.0064), with significant differences between original-no decontamination and water-no decontamination (Tukey’s HSD, p=0.0026 and p=0.0206, respectively). Finally, for enamelin, no significant differences between methods in terms of covered positions were found (ANOVA, F=2.559, p=0.0734). Summarized across the three investigated enamel proteins, the highest number of covered amino acid positions was found in the original samples (326.0±47.09) and the lowest in the fully contaminated samples (215.33±17.01), with significant differences between the methods (ANOVA, F=4.818, p=0.0085). Significant between-method differences were found between original-no decontamination and water-no decontamination (Tukey’s HSD, p=0.0036 and p=0.0284, respectively).

**Figure 5.**
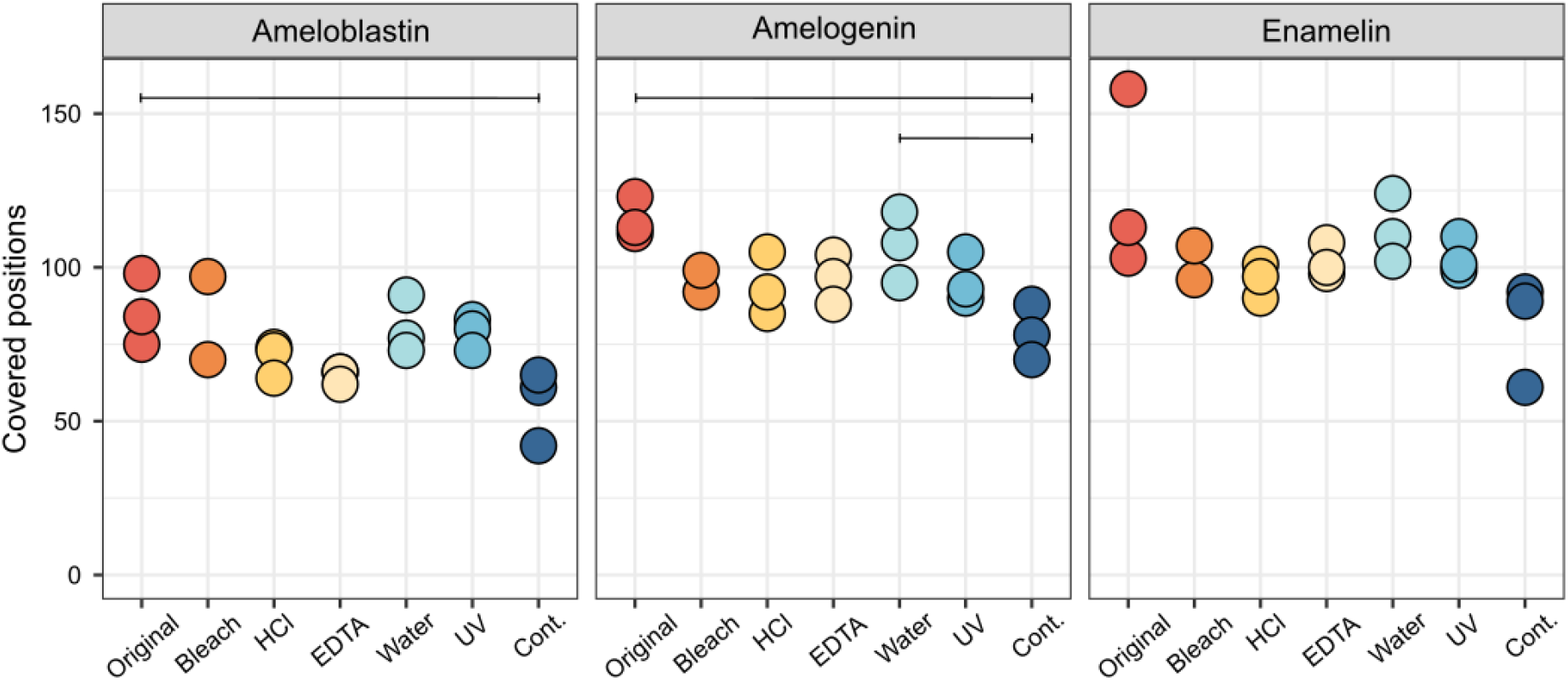
Number of covered positions of ameloblastin (AMBN), amelogenin (AMEL*) and enamelin (ENAM) after different decontamination methods for each replicate. Horizontal lines indicate statistically significant differences between methods. Cont. = Contaminated.

Mean length of recovered enamel peptides is significantly different between the decontamination methods (ANOVA, F=3.679, p=0.0233; Figure 3b). Significant between-treatment differences are only present between original-bleach (Tukey’s HSD, p=0.0490), with the bleach-washed samples having longer peptides. It should, however, be noted that this is only a change of mean length from 12.76±0.09 amino acids for original samples, to 13.27±0.18 amino acids for bleach-treated samples. In part, this result might be expected given that bleach treatment is known to remove intercrystalline protein fragments, which are expected to be more degraded, and retain only intracrystalline protein fragments, which can be expected to be less degraded ^22^. This would, on average, increase mean peptide length. Mean peptide hydrophobicity is also significantly different depending on the used method (ANOVA, F=14.36, p<0.001). These differences are significant between bleach-EDTA, bleach-HCl, bleach-water, bleach-UV and bleach-no decontamination, as well as between original-EDTA, original-HCl and original-no decontamination, between water-no decontamination, and between UV-no decontamination (Tukey’s HSD, p<0.05 in each case).

The fraction of phosphorylated serines is found to differ significantly depending on the decontamination method (ANOVA, F=7.012, p=0.0017), with significant differences between original-HCl, original-no decontamination, as well as no decontamination-UV, and no decontamination-water (Tukey’s HSD, p<0.05 in each case). This indicates that the amount of peptides containing phosphorylation is brought back to normal levels by UV irradiation and a water wash. None of the methods significantly affect deamidation of N or Q (ANOVA, F=2.227, p=0.0582 and F=1.02, p=0.454, respectively). Taken together, these analyses of peptide and amino acid properties indicate that the retrieval of enamel-derived proteomic information is mainly improved through a water wash, as there are significant differences between fully contaminated and water-washed samples.

## Discussion

Archaeological materials analysed through biomolecular methods often go through a long journey from the archaeological site to the laboratory, a process that sometimes encompasses decades. During this time, the objects will have accumulated modern contamination, through different types of handling, study, transport and storage. This leads to extracted biomolecular assemblages consisting of a mixture of modern and ancient, exogenous and endogenous molecular information, unless the contamination is removed. In the case of the palaeoproteomic study of dental enamel, it has often been assumed that protein decontamination is unnecessary due to the nature of the enamel proteome. First, most enamel proteins are unique and not expressed elsewhere in the body, leading to, presumed, straightforward bioinformatic separation of recovered proteins into contaminants and endogenous molecules. Second, as enamel proteins are digested *in vivo*, palaeoproteomic extraction protocols for enamel often do not contain enzymatic digestion steps^1^, and thus modern contaminant proteins would be too intact to be successfully analysed through mass spectrometry. These assumptions about protein contamination have, however, not been empirically tested.

Through artificial contamination of enamel from a Pleistocene woolly rhinoceros, we show that contamination can indeed affect palaeoproteomic dental enamel analyses negatively, although contaminating proteins are not identifiable. These negative effects are mainly visible as a loss of proteomic information, and a slight bias in peptide recovery. Low-abundance proteins and protein regions may be masked, due to the abundance of the contaminating molecules hindering acquisition of mass spectra from enamel proteins. If these regions contain polymorphisms that are essential for differentiation between taxa, phylogenetic studies may not be successful without proper protein decontamination. The effects may be even more severe for very ancient specimens, with low amounts of preserved endogenous proteins. The artificial protein contamination experiment has allowed us for the first time to demonstrate the effects of protein contamination on palaeoproteomic analyses of dental enamel, indicating the necessity of decontamination to ensure that a maximal amount of endogenous proteomic information is gained from each sample.

Comparison of five different published decontamination methods shows varying success at restoring protein recovery to pre-contamination levels. Of the tested methods (wash with bleach, HCl, EDTA, and water, as well as UV irradiation), we find that washing the enamel sample with molecular grade water shows the best results. This is different to what was found for Pleistocene bone, where a bleach wash was most successful^13^. The difference may be due to the differing mineral composition and density between bone and dental enamel^23^, leading to the contaminating proteins binding less strongly to the dental enamel. As a water wash is unlikely to interfere with downstream protein extraction, this is a very short and simple step to add to established protein extraction protocols, to improve protein recovery from dental enamel. It should be noted, however, that as the bleach treatment likely removes intercrystalline proteins and free amino acids^22^, some of the metrics utilized here to evaluate decontamination success, such as the number of identified MS2 spectra and peptide length, are difficult to directly compare, and a bleach wash may be more successful than the explored metrics show.

Thus far, two different archaeological materials have been compared with regard to palaeoproteomic decontamination protocols: bone^13^ and dental enamel (present study). Despite using the same artificial contamination method, different methods were found to be optimal (bleach vs. water). This shows the necessity of experimental decontamination studies for various archaeological materials. Several non-skeletal materials are also commonly studied palaeoproteomically, such as dental calculus, leather, parchment and ceramics^10^. It can be expected that these require different decontamination approaches in order to efficiently remove contaminating proteins without damaging the endogenous proteome. Additionally, samples of different preservation states may require optimized approaches, as the material degrades over time.

A common result for both bone and dental enamel is, however, that decontamination is necessary in order to avoid loss of proteomic information. Contamination has been shown to mask endogenous proteins, hindering the recovery of low-abundance proteins and protein regions. In studies targeting very abundant proteins, such as taxonomic studies utilizing collagen type I, this may be less of an issue. However, in studies targeting low-abundance protein ^24^ or aiming to reconstruct as much of the proteome as possible for phylogenetic studies^5,13,25,26^, loss of low-abundance peptides can have a significant negative effect. It is therefore recommended to decontaminate archaeological and palaeontological samples prior to protein extraction, using a method appropriate for the material in questions, to ensure that the maximal amount of endogenous information is acquired from each destructive sampling.

Decontamination has previously been shown to be a complex question within archaeological sciences. For example, for ancient bone, bleach decontamination has been found to be highly efficient at removing contamination, albeit at the cost of a loss of a significant amount of endogenous DNA^27,28^. More gentle methods, such as a pre-digestion with EDTA and proteinase K^29^ or phosphate buffer^27,30^ are less efficient at removing contamination, but may be more suitable for highly degraded specimens to avoid complete destruction of endogenous DNA. Bleach decontamination is also commonly employed for amino acid racemization (AAR) analyses, as it retains only intracrystalline amino acids^22^. We hypothesize that the loss of intercrystalline proteins through bleach treatment can also be seen in the present study, through a lower amount of recovered endogenous proteomic information, while simultaneously resulting in the recovery and identification of longer peptides. This indicates that although a bleach treatment may be destructive to a portion of the endogenous proteins, namely those present in the intercrystalline compartment, it simultaneously retains the better-preserved intracrystalline proteins. The observations made on our bleach-treated dental enamel samples are therefore in line with the expectations derived from AAR analysis on dental enamel^22^.

The issue of protein contamination detection and removal in palaeoproteomics is complex and requires thorough study to identify appropriate methods. It is a trade-off between efficiency of contaminant removal and destruction of endogenous proteins. We have now demonstrated that protein decontamination is, however, essential for both bone and dental enamel prior to protein extraction, since its unrecognized presence will mask the identification of low-abundance, endogenous proteins and peptides. Furthermore, for both bone and dental enamel, to avoid loss of endogenous protein information from archaeological and palaeontological materials, chemical extraction procedures are now available that significantly remove exogenous protein contamination while avoiding the destruction or chemical modification of endogenous peptides. These methodological advances will enable the study of highly contaminated archaeological and palaeontological specimens.

## Methods

### Specimen

Visible dentine was removed by drilling from a tooth fragment from a Pleistocene woolly rhinoceros (*Coelodonta antiquitatis*) from the Zandmotor, the Netherlands. Based on its taxonomic identity and the composition of fossils found on the Zandmotor artificial sand bank, the specimen can be expected to be Late Pleistocene in chronological age^31^. Part of the resulting dental enamel fragment was powdered by mortar and pestle. The rest of the fragment was artificially contaminated by saliva from a dog, who had not eaten recently prior to the contamination and had no known oral disease at the time. Dog saliva was chosen for the contamination based on it being a large, complex proteome^32^ which absorbs to skeletal elements, and has previously been successfully used to artificially contaminate Pleistocene bone^13^. The contaminated enamel fragment was crushed into a rough powder by mortar and pestle and divided into the different tested decontamination approaches.

### Protein extraction

On average 11.1 mg (range 9.9-12.5 mg) of enamel powder was used for each protein extraction (Supplementary Data 1). Proteins were extracted from the original (uncontaminated) enamel and the contaminated (non-decontaminated) enamel, to provide baselines for original and fully contaminated proteomes. Five different methods for decontamination were tested, following Fagernäs et al. 2025^13^: 0.5% bleach (sodium hypochlorite), 1% HCI (hydrochloric acid), 5 M EDTA (ethylenediaminetetraacetic acid), molecular grade water, and UV irradiation (UVC at 254 nm). The percentage of HCl used was lower than in Fagernäs et al. 2025^13^ to avoid demineralization of the enamel powder during the decontamination step, which preliminary tests showed to be an issue. Each method was tested in triplicate. For all wash methods, 1 ml of each reagent were added to their respective samples, whereafter they were vortexed for 5 seconds, centrifuged for 1 minute at 13,000 rpm, and the supernatant was removed. The remaining pellets were washed twice with 0.5 ml molecular grade water after decontamination with bleach and EDTA, in order to avoid the decontamination reagents interfering with downstream protein extraction. UV irradiation was conducted by keeping the enamel powder in open tubes, irradiating for 30 s in a crosslinker (UVP Crosslinker, Analytik Jena), whereafter the tubes were lightly shaken and the irradiation repeated (total irradiation time 1 min).

For all samples, including the original and non-decontaminated samples, a standard dental enamel palaeoproteomic extraction protocol was followed^1,2^. The pellets were suspended in 1 ml of 5% HCI and demineralized overnight at 4°C under agitation. The samples were then centrifuged for 10 min at 13,000 rpm, whereafter the supernatant (S1) was removed and stored at −20°C. A fresh 1 ml 5% HCI was added to all samples in order to complete the demineralisation overnight at 4°C under agitation. The supernatant was removed after centrifuging for 10 minutes at 13,000 rpm (S2). Proteins were desalted using in-house made Stagetips following Cappellini et al. 2019^2^. Briefly, 150 μl of methanol was centrifuged through the Stagetips, followed by 150 μl of AT80 (80% of 0.1% TFA in ACN and 20% of 0.1% TFA in H_2_O). Thereafter, 150 μl of 0.1% TFA in H_2_O was added to the Stagetips and centrifuged through. The supernatants (S1 and S2) of the samples were then centrifuged through, and the Stagetips were cleaned twice with 0.1% TFA in H_2_O and stored at −20°C until LC-MS/MS analysis.

### LC-MS/MS

Mass spectrometry was carried out at the Centre for Protein Research at the University of Copenhagen (Denmark). Liquid chromatography was carried out on a Vanquish Neo UHPLC system (Thermo Fisher Scientific, Waltham, MA, USA). Peptide eluate was loaded at 600 bar pressure. Separation was carried out at a flow rate of 250 nl/min at a temperature of 275 °C. Mass spectrometry was performed on an Exploris 480 (Thermo Fisher Scientific, Waltham, MA, USA) at 2 kV positive ionization mode. The ion transfer tube was maintained at 275 °C. The MS1 scan was from 350 to 1400 m/z with an orbitrap resolution of 120,000 and the MS2 experiment with m/z from 100 to precursor m/z + 20 at a resolution of 60,000. An HCD collision energy of 30% was used for fragmentation. The top 10 ions with an intensity above 2e4 with charge state 2-6 were fragmented, and then excluded for 20s. The raw data have been deposited to the ProteomeXchange Consortium via the PRIDE^33^ partner repository with the dataset identifier PXD064417.

### Analysis

The raw data was analysed using MaxQuant v. 2.1.3.0^21^ with a database consisting of enamel proteins^1,5^, bone proteins^34^, and salivary proteins identified in a previous decontamination experiment^13^. This resulted in a total of 389 proteins that were included (Supplementary Data 2) from both the south-central black rhinoceros (*Diceros bicornis minor*) or, when unavailable, the southern white rhinoceros (*Ceratotherium simum simum*), as well as dog (*Canis lupus familiaris*). The rhinoceros species were chosen based on the number of available published protein sequences. Rhinoceros sequences were downloaded from NCBI on 2024-03-13 and dog sequences from the reference proteome (UP000805418) from Uniprot on 2022-06-10. It should be noted that no AMELY sequences for Rhinocerotidae were available and therefore any analysis of amelogenin is based solely on peptides matching AMELX. The internal MaxQuant contaminant database was also used, as it contains common laboratory contaminants, as well as human keratins, which may have been present on the sample prior to the artificial contamination. Unspecific digestion was used, and deamidation (NQ), oxidation (P) and phosphorylation (STY) were set as variable modifications.

Statistical analyses were conducted in R v.4.5.0^35^ using a one-way ANOVA, with input variables scaled by weight of enamel used for extraction. Homogeneity of variances was checked using Levene’s Test, and normality of residuals using a Shapiro-Wilk Normality Test. R packages *MetBrewer* v.0.2.0^36^, *janitor* v.2.2.1^37^, *ggpubr* v.0.6.0^38^, *car* v.3.1.2^39^, *Peptides* v.2.4.6^40^, *Hmisc* v.5.2.3^41^ and *tidyverse* v.2.0.0^42^ were used for analyses and visualization. Peptide numbers are based on the razor+unique count from MaxQuant. During preliminary data analysis, it was noted that one of the bleach-replicates represents a clear outlier with regard to enamel protein recovery, as it has 84 enamel PSMs, whereas the two other replicates have 378 and 466 PSMs, respectively. It is therefore excluded from all analyses specifically focusing on enamel proteins. R code used for analysis has been archived on Zenodo at 10.5281/zenodo.15553261.

## Acknowledgements

This research has been made possible through funding from the European Research Council (ERC) under the European Union’s Horizon 2020 research and innovation programme, grant agreement no. 948365 (PROSPER, awarded to F.W.), the European Union’s Horizon Europe research and innovation programme under the Marie Skłodowska-Curie grant agreement no. 101106627 (PROMISE, awarded to Z.F.), and through generous funding from the Leakey Foundation (awarded to Z.F.). Views and opinions expressed are those of the author(s) only and do not necessarily reflect those of the European Union or the European Research Council Executive Agency. Work at the Novo Nordisk Foundation Center for Protein Research is funded in part by a donation from the Novo Nordisk Foundation (grant number NNF14CC0001). Finally, we thank the Contaminator, Tjorven (Tastaway’s Herrmann), for his contribution to the research.

## Author contributions

Z.F. and F.W. designed the research. J.K.B. contributed archaeological samples. J.V.O., and F.W. ensured access to the required equipment. Z.F., S.S. and G.T. performed laboratory research. Z.F. analyzed the data. Z.F. and F.W. prepared the manuscript, with input from all coauthors.

## Data availability

The mass spectrometry proteomics data have been deposited to the ProteomeXchange Consortium via the PRIDE^33^ partner repository with the dataset identifier PXD064417. R code used for analysis has been archived on Zenodo at 10.5281/zenodo.15553261.

